# A *Dynamic Affective Core* to bind the Contents, Context, and Value of Conscious Experience

**DOI:** 10.1101/2021.05.20.444839

**Authors:** Kenneth T. Kishida, L. Paul Sands

## Abstract

The private and dynamic nature of conscious subjective experience poses an empirical challenge that has led neuroscience-based theories about consciousness to note the importance of ‘the hard problem’ of explaining how subjective phenomenal experience can arise from neural activity but set it aside and focus on the ‘easier’ problems associated with information representation and behavior. This approach leaves a major gap in our understanding of the neural mechanisms underlying conscious subjective experience and its dynamic nature. However, computational methods integrated with a variety of tools for measuring human brain activity are beginning to link dynamic changes in subjective affect with reproducible neurobehavioral signals in humans. In particular, research applying computational reinforcement learning theory has shown tremendous utility in investigating human choice behavior and the role the dopaminergic system plays in dynamic behavioral control. This research is beginning to reveal an explicit connection between the dynamics of dopaminergic signals and dynamic changes in subjective affect. However, it should be obvious that the dopaminergic system alone is not sufficient to explain all of the complexities of affective dynamics. We review foundational work, highlight current problems and open questions, and propose a *Dynamic Affective Core Hypothesis* that integrates advances in our understanding of the representation of the content and context of conscious experiences with our nascent understanding about how these representations acquire and retain affective subjective value.

## Introduction

In “What is it like to be bat?” (Nagel 1974), Nagel highlights the gap in our ability to provide a mechanistic account of subjective phenomenal experience from our current knowledge about nervous systems. In a related vein, Chalmers (1996) coined the distinction between ‘easy’ problems and ‘the hard problem’ facing investigations about how physical processes generate subjective phenomenal experience (i.e., *qualia*). The ‘*easy problems*’ are those for which it is conceivable that we will find solutions given the currently known mechanistic working of neural processes; ‘*the hard problem’* concerns an explanation of how the physical processes could possibly give rise to subjective phenomenal experiences *a priori*. Leading neuroscientific theories about consciousness note the importance of subjective phenomenal experience but set this ‘hard problem’ aside to instead focus on the 'easy problems' regarding how nervous systems represent information and control behavior (Edelman and Tononi, 2001; Crick and Koch, 2003). In line with others’ ‘faith’ in a scientific approach (Churchland and Churchland, 2002; Churchland, 2005), we reject this distinction. 'The hard problem' is hard, but not in any special way that prevents scientific investigation. Instead, it represents the most exciting frontier in human neuroscience research. Before we had an empirically based theory of electromagnetism, electricity and magnetism must have seemed magical and fundamentally unexplainable (through known physical mechanisms) to philosophers of the time. But, through rigorous empirical investigation and the development of supporting mathematical theory, major breakthroughs came in our understanding of a fundamental physical phenomenon (Forbes and Mahon, 2014). We believe that (broadly) neuroscience research has simply not focused its attention, until recently, to the problem of conscious subjective experience in humans. These tides are changing, and advances in seemingly disparate areas of research are poised to come together through applications of mathematical theory to begin to shed light on how a nervous system could give rise to subjective phenomenal feeling.

‘Affective dynamics’ – and particularly computational approaches thereof (Cunningham et al., 2013) – represent an investigative construct well-suited to push past the current boundaries of neuroscience research about dynamic changes in emotion-related behavior and move towards a neuroscience of consciousness squarely focused on mechanisms giving rise to subjective phenomenal experience. Presently, much of the work can be framed into two (admittedly overbroad) domains of research: 1) investigations into the role of reinforcement learning in driving dynamic changes in behavior and associated reports about subjective emotional reactions, and 2) investigations into the roles of functional networks involving cortical and sub-cortical brain regions in dynamically representing emotion. While both domains of research are in and of themselves rich and highly productive, little has been done to bridge the divide between them and provide a mechanistic account of the dynamics underlying how changes in the environment or behavioral state of an individual induce changes in the functional networks representing the vast range of observable, dynamic emotional reactions and associated subjective experiences.

Here, we discuss a framework for investigating affective dynamics that is founded on computational reinforcement learning theory (Sutton and Barto, 1998) and its tight connection to the neurobiology of adaptive behavior. We review the dominant theory for dopamine neuron function (as a generator of a “temporal difference reward prediction error” signal) and its role in behavioral control and value updating (Montague et al., 1996; Schultz et al., 1997; Montague et al., 2004; Glimcher, 2011; Watabe-Uchida et al., 2017). We review how this framework has been used to date to investigate affective dynamics, but also this approach’s dopamine-centric limitations. We follow this discussion with an extension of temporal difference reinforcement learning theory to include a parallel system hypothesized to support *temporal difference aversive learning*, 2) the interaction of these parallel appetitive-learning and aversive-learning systems to generate valuation and salience signals, and 3) evidence that these valuation and salience signals are fundamental to dynamic affective experience. We briefly discuss a candidate set of key neural structures that we hypothesize are necessary components of a dynamic network for representing emotion, and we describe how ascending valuation systems (e.g., dopamine- and serotonin-releasing neurons) are integral to this network. We discuss how this network relates to the *dynamic core* hypothesis that was proposed by Edelman and Tononi (2001) to explain consciousness – in particular, we note that their notion of a *dynamic core* did not necessitate key elements, and the consequences of these omissions. Crucially, we believe that Edelman and Tononi’s ‘dynamic core’ appears sufficient for an integrated representation of the contents and context of dynamically evolving conscious experiences, but it omits systems required to provide those representations affect and value. Thus, we introduce the *Dynamic Affective Core* hypothesis, which updates the original dynamic core hypothesis to now necessarily include originally omitted affect and valuation systems. We discuss the implications of the Dynamic Affective Core hypothesis and future directions for this line of research.

## Affective Dynamics as a phenomenon resulting from systems seeking ‘Optimal Control’

Moment-to-moment changes in one’s emotional state can be driven by external and internal signals. The environment naturally evolves, and an organism’s nervous system ought to track this evolution with its own evolving representations of the environment in parallel with representations of its own evolving internal states (e.g., proprioceptive body position, body temperature, energy levels, etc.). Associated with these objective quantities are affective, subjective feelings (e.g., hunger, thirst, pain, and pleasure). These representations are used in turn to guide adaptive physiological responses and behavior. How nervous systems accomplish this can be investigated through a computational framework (Minsky, 1967) where the actual “state of an agent” and the actual “state of the environment” can be represented explicitly, as can their interactions. Further, *representations* of the state of an agent and the environment can be explicitly represented as hypothetical physical states that nervous systems may instantiate. Within such a framework, one can consider that as the state of the agent and external environment continuously evolve, the affective state of the agent is expected to also continuously evolve. We hypothesize that the representation of exteroceptive and interoceptive state-transitions is associated with representations of the affective emotional valence (i.e., subjective value) of these experienced states and actions. This dynamic evolution of emotional subjective experience is the target of our investigation. To be rigorous in our approach, we employ a computational framework to aid in making explicit our assumptions and hypotheses about the dynamic mechanisms at play.

Artificial intelligence research has developed theory around how the behavioral problem (e.g. how to choose how to act and adapt behavior) may be solved with an eye towards optimal solutions. One particularly successful line of reasoning has led to the development of computational reinforcement learning theory (Sutton and Barto, 1998 and 2018). Within artificial intelligence research, computational reinforcement learning theory has been used as a core theoretical construct to explore how emotions may be integrated into the decision-making processes of computational (artificially intelligent) agents (surveyed in Moerland et al, 2018). More specific to our line of inquiry is recent empirical work bridging computational reinforcement learning theory and human neuroscience that is beginning to connect dopamine neuron activity with how the human brain dynamically adapts not only representations of states and actions, but also representations of associated subjective experiences (Xiang et al., 2013; Rutledge et al., 2014; Eldar and Niv 2015).

### Computational Reinforcement Learning Theory and Dopamine

Computational Reinforcement Learning (RL) theory (Sutton and Barto, 1998, 2018) provides an explicit framework to investigate processes involved when a theoretical agent makes decisions under uncertainty, experiences the consequences of those decisions, and makes changes in its approach (i.e., learns) to make ‘better’ decisions in the future. This theoretical framework assumes that the *agent* seeks to maximize the ‘reward’ it attains and builds from this fundamental assumption a mathematical rendering that incorporates theory about Finite Markov Decision Processes, Dynamic Programming, and Monte Carlo Methods (see Sutton and Barto, 1998 and a revised edition in 2018 for an authoritative text book on how these ideas are brought together). Importantly, the agent in question is an abstract computational agent – not necessarily a biological one, let alone human. This approach simply seeks to find computable optimal solutions to behavioral control and, when such computations are intractable, to determine best estimates for these solutions (Bach et al., 2017). Moerland and colleagues (Moerland et al., 2018) have recently provided an extensive survey of how these ideas have been explored to incorporate models of ‘emotions’ into various learning algorithms in AI research. Moerland’s survey provides a wide range of hypotheses about how emotions may be represented in adaptive reinforcement learning algorithms, but empirical support that these proposals are actually implemented biologically remains lacking. On the other hand, Montague and colleagues’ publication in 1996 (Montague et al, 1996) demonstrated that dopamine neurons in non-human primates encode a key signal in reinforcement learning theory – a *temporal difference reward prediction error*. This finding has since been a foundation for research into the basis of adaptive decision-making in humans (and non-human animals) and more recently in investigations about the neural basis of dynamic changes in subjective experience in humans.

*Temporal Difference Reinforcement Learning* is a foundational idea that grew out of computational reinforcement learning theory research. The basic idea is as follows. It is assumed that an agent always seeks to maximize the attainment of ‘reward’. To do so, it must learn the value of various states and actions that it finds itself in and that it has available to it at any given time, respectively. The ‘Value’ (*V*_*t*_) of a particular state at a given time (*t*), in this context, is simply the sum of the reward currently acquired (*r*_*t*_) plus the rewards the agent may attain in the future given that it finds itself in that state (*r*_*t+1*_ + *r*_*t+2*_ + *r*_*t+3*_ + ···), so:

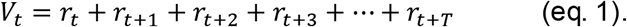

We can modify this estimate by incorporating a discounting of future (not yet acquired) rewards:

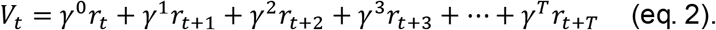

Here, *γ* is a parameter that is greater than zero, less than one, and therefore shrinks exponentially small when raised to increased powers towards *T*. This causes proximate rewards to be weighted more heavily than distant future rewards. The challenge for the agent (with incomplete knowledge of the future) is to estimate this value function for any given state it may find itself in. Learning (better estimates of *V*_*t*_) proceeds through experience (real or simulated) where prediction *errors (*δ) about the estimated expected value are used to update (i.e., make corrections to) the estimated expected value:

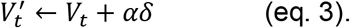

In equation 3, *V*_*t*_ on the right-hand side is changed to 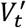 by an amount *δ* (i.e., the reward prediction error) modulated by the learning rate *α*. Small fractional changes in *α* lead to slower learning, whereas larger *α* causes *δ* to have a bigger influence. In *temporal difference reinforcement learning*, the errors – called *temporal difference reward prediction errors* (*δ*) – are calculated a s follows:

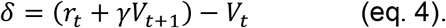

Note that these errors are determined by incorporating not only the experienced reward in the current state (at time “t”: *r*_*t*_), but also a discounted estimate of future rewards (*γV*_*t*+1_); combined, these two terms ((*r*_*t*_ + *γV*_*t*+1_)) are compared to the current estimate of value (*V*_*t*_) and the difference is the “temporal reward prediction error”. This reward prediction error term has been hypothesized and demonstrated to be encoded by dopamine neuron activity in non-human primates (Figure 1: Montague et al., 1996; Schultz et al., 1997; Bayer and Glimcher, 2005; reviewed in Glimcher 2011; for a historical account: Colombo, 2014; Watabe-Uchida et al., 2017) and to be consistent with temporal dynamics of blood oxygen level dependent responses (measured with functional magnetic resonance imaging) in regions of the human brain that are highly innervated by dopamine releasing terminals (Pagnoni et al., 2002; Montague et al., 2006).

**Figure 1.**
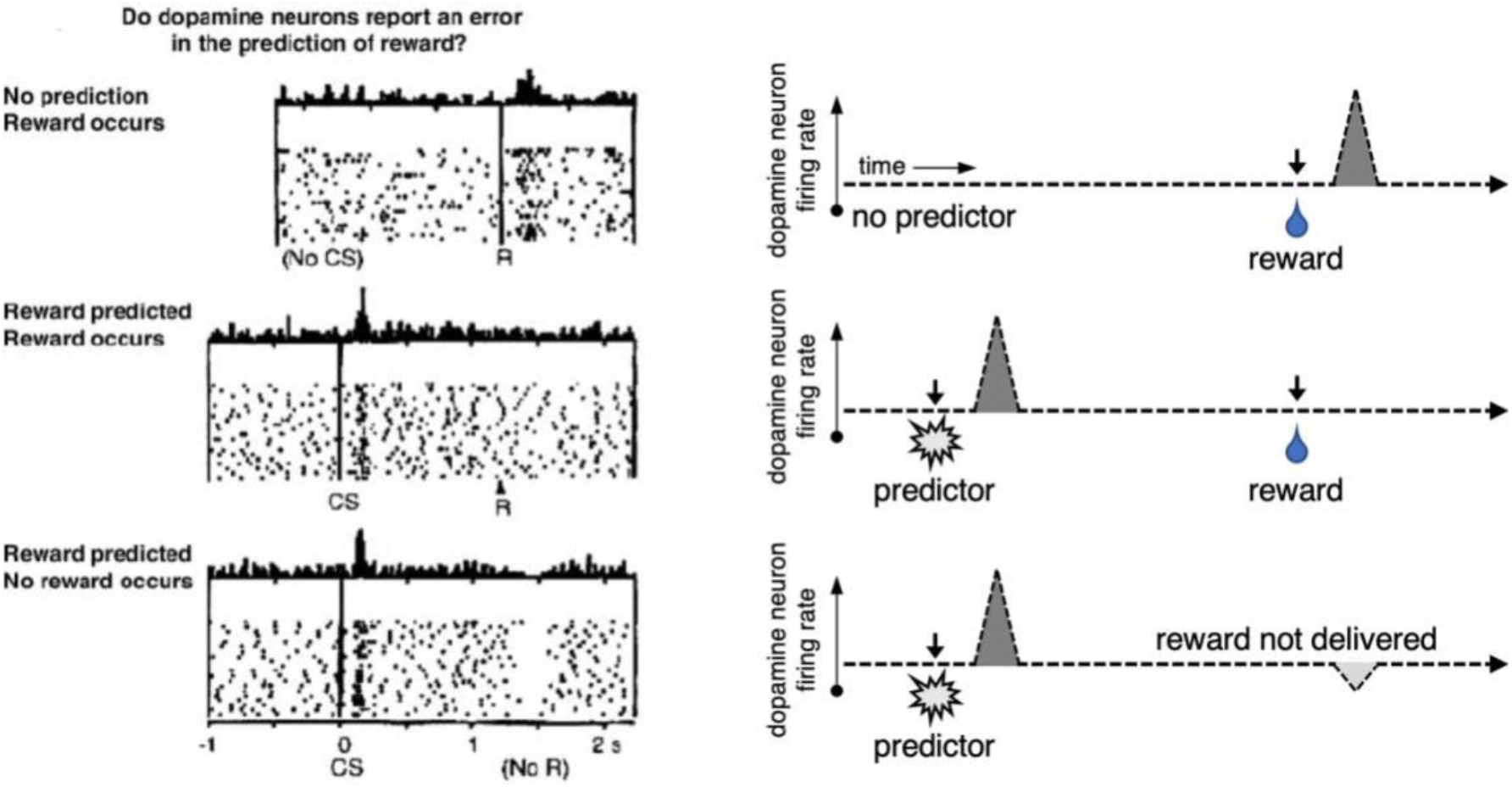
Dopamine neurons encode temporal difference reward prediction errors. Dopamine neurons change their rate of firing in a manner consistent with ‘temporal difference reward prediction errors’. Left: Recordings of dopamine neuron spike rates during the presentation of rewards (squirts of juice) and a conditioning stimulus (e.g. flash of light); figure panel adapted from Schultz, Dayan, and Montague, 1997. Right: Cartoon depiction of dopamine neuron behavior in the different phases of learning depicted in left panel. Top row: Prior to learning, dopamine neurons increase their firing rate in response to unexpected delivery of reward. Middle row: After conditioning, with consistent pairing of a predictive stimulus and a reward, dopamine neurons increase their firing rate to the predictive stimulus and do not change their firing rate when the reward is delivered as expected. Bottom row: After conditioning, dopamine neurons increase their firing rate to the predictive stimulus and go silent (firing rate goes to zero) when the expected reward is not delivered.

The first association between the temporal difference reward prediction error and non-human primate dopamine neuron activity was shown in Montague et al., 1996 and reviewed in the context of prior work in Schultz et al., 1997. Subsequently, the role reward prediction errors and dopaminergic signaling plays in mammalian decision making grew into a highly impactful field of research (reviews: Montague et al., 2004; Glimcher 2011; Colombo, 2014; Watabe-Uchida et al., 2017), with elaborations on this basic framework revealing a consistent role for reward prediction error signals guiding human and animal behavior during a wide variety of experimental paradigms that require these agents to make incentivized decisions under uncertain conditions.

A number of predictions from this basic formulation have been tested and the overarching hypothesis upheld, though recent work has challenged the completeness of this idea in humans (Kishida et al., 2011; Kishida et al., 2016; Moran et al., 2018; Bang et al, 2020). The *“temporal difference reward prediction error”* (eq. 4) *hypothesis* for dopamine neuron activity, expressed in words predicts: (1) increases in DA neuron spike activity (from background rates) when “things are better than expected”, (2) decreases when “things are worse than expected”, and (3) no change when “things are just as expected”. In this interpretation, dopamine neurons are always emitting information to downstream neural structures since even no change in firing rate carries meaning. This computational hypothesis is strongly supported by the *timing* and *amplitude* of burst and pause responses in the spike trains of dopamine neurons (Montague et al., 1996; Schultz et al., 1997; Fiorillo et al., 2003; Montague et al., 2004; Bayer and Glimcher, 2005; Dayan and Niv, 2008; Watabe-Uchida et al., 2017). Further, this hypothesis appears to be supported (at least in part) by dopamine release measurements in rodents (Hart et al., 2014, Clark et al., 2010), single unit recordings of dopamine neurons in humans (Zaghloul et al., 2009), and direct measurements of dopamine release in humans (Kishida et al. 2016; Moran et al., 2018).

Note, however, that in Kishida et al., 2016; and Moran et al. 2018 it was demonstrated that sub-second changes in dopamine levels in human striatum are not fully explained by the simple reward prediction error hypothesis. In these experiments, extracellular dopamine release was measured in humans with sub-second temporal resolution during a sequential monetarily incentivized decision-making task (Kishida et al., 2011; Kishida et al., 2016). While these measurements were made, participants performed a stock market gambling task that elicited reward prediction errors about the participants’ investment decisions and market fluctuations. Importantly, participants’ investments were lodged as percentages of their continuously updated portfolio. Thus, counterfactual information – what “could have been” had they chosen to invest more or less than they actually did – was present on every trial. Kishida et al., 2016 demonstrated that sub-second dopamine fluctuations in the striatum integrated the reward prediction error term with a counterfactual prediction error term (Figure 2). In other words, dopamine fluctuations in response to better- or worse-than-expected outcomes (i.e., reward prediction errors) were depressed or even inverted according to the magnitude of missed gains or avoided losses had the participants invested one-hundred percent of their portfolio. In this sense, measured fluctuations in dopamine levels within ~700msec following an outcome corresponded with how participants *ought* to have felt: *positive feelings about better-than-expected outcomes* would be muted or even inverted to negative feelings when regret (over not investing more) increases, and *negative feelings about worse-than-expected outcomes* would be muted or inverted to positive feelings when relief is high (Figure 2, ‘regret’ and ‘relief’ would increase with decreasing bet size). In a follow up study, Moran and colleagues extended the work and demonstrated that serotonin fluctuations in human striatum encoded an opponent response to the dopamine signal, suggesting that dopamine and serotonin are, together, critical in the processing of actual and counterfactual reward prediction errors in sequential decision-making processes. Unfortunately, however, subjective assessments were not performed in these experiments, so the connection between sub-second dopamine and serotonin release and subjective experience remains hypothetical.

**Figure 2.**
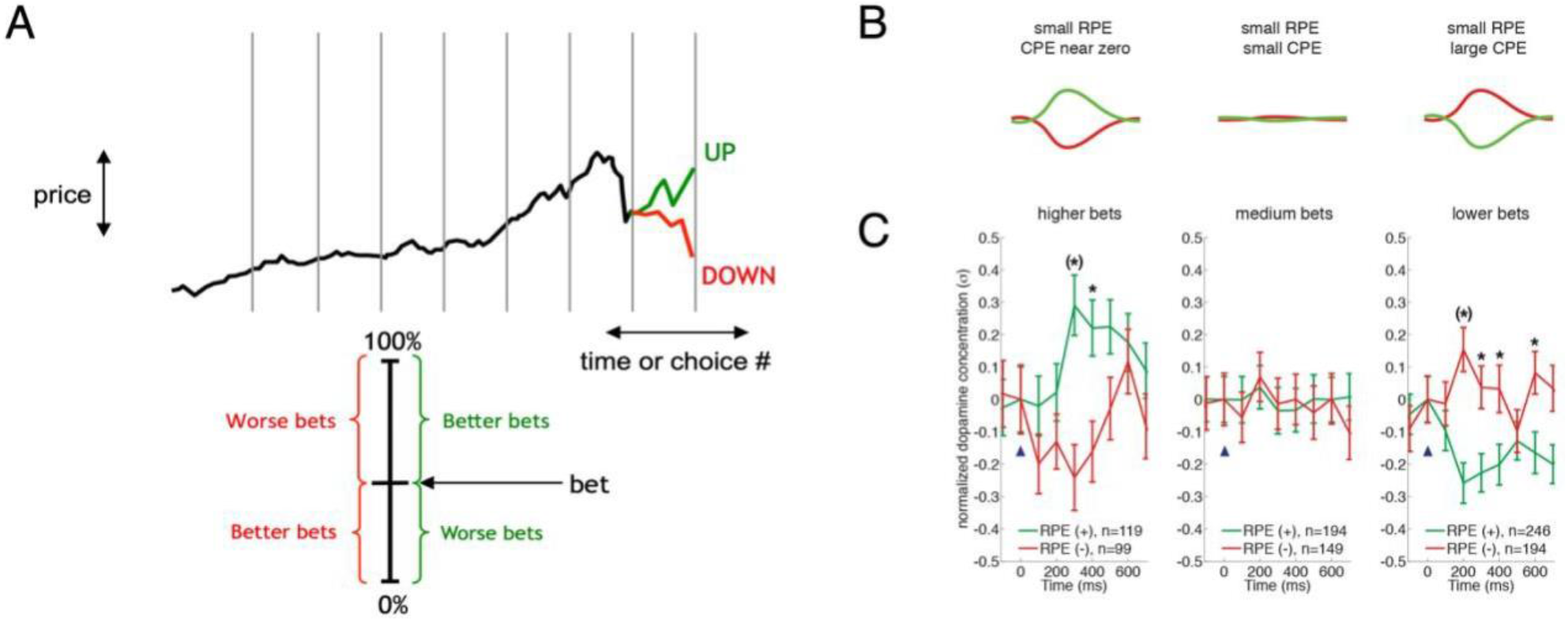
Dopamine release in human striatum encodes superposed *reward prediction errors* and *counterfactual prediction errors*. Direct recordings of dopamine release during a sequential decision-making task (A) demonstrates that dopamine release encodes reward reward prediction errors (left columns of B & C), but that counterfactual information diminishes or inverts dopaminergic encoding of reward prediction errors about actual experience (middle and right columns of B & C). Figure panels adapted from Kishida et al., 2016. **A.** Sequential investment game: Participants ‘invest’ into a stock market by lodging bets as a percentage of their portfolio. Market returns increase or decrease the participant’s portfolio as a function of the percent invested and the percent change in the stock price. For market increases and decreases, the amount not invested represent missed gains or avoided losses, respectively. These counterfactual outcomes are observed to modulate dopamine (panel C) and serotonin (Moran et al., 2018) responses to reward prediction errors over actual gains and losses in the game. **B.** and **C.** Model predictions (B) and actual dopamine responses (C) for a model that integrates ‘reward prediction errors’ and ‘counterfactual prediction errors’. Green: dopamine responses (predicted (B) or observed(C)) to positive reward prediction errors. Red: dopamine responses (predicted (B) or observed(C)) to negative reward prediction errors. Left column: Counterfactual outcomes are minimized (investments near 100%) and the dopamine response is positive for positive reward prediction errors and negative for negative reward prediction errors (as predicted by the traditional reward prediction error hypothesis). Middle and Right columns: As the bet sizes decrease, the counterfactual terms increase and diminish (middle column) or invert (right column) the dopamine response to positive and negative reward prediction errors. Figure panel C adapted from Kishida et al., 2016.

The hypothesis that dopamine fluctuations encode fluctuations in positive affect may seem obvious, given that dopamine neurons and extracellular dopamine levels dynamically encode “reward” prediction errors. This is consistent with pleasure and subjective well-being focused emotion literature connecting these emotions and related processes to dopamine’s function (Kringlebach and Berridge, 2017). However, it is important to note that the term “reward” in the naming of this error term does not *explicitly* mean the “subjective experience of reward”. Rather, this term refers to a quantity that a computed objective function aims to maximize, and its connection to subjective reward is only implied or useful as an analogy when thinking about related computer algorithms from an anthropomorphic perspective. The explicit relationship between reward prediction errors, dopamine, and dynamic changes in human subjective experience has only recently begun to be explored.

### Reinforcement Learning and Affective Dynamics

The formal framework of computational reinforcement learning provides a rich landscape for investigating how the brain may generate dynamic changes in emotional behavior and subjective experience (Doya 2000; Xiang et al., 2013; Eldar et al, 2016; Bach and Dayan, 2017; Huys and Renz, 2017; Moerland et al., 2018). For example, the development and implementation of computer agents that use modified reinforcement learning algorithms to incorporate intuited features of emotion can generate a wide spectrum of hypotheses about how the human brain may ‘compute’ and generate emotional behaviors with the goal of augmenting artificial intelligence behavior (Moerland et al., 2018). While this approach notes the connection between reinforcement learning and hypothesized neural processes, their primary focus on machine learning applications is less constrained by or interested in solving the biological question and more concerned with the augmentation of machine learning algorithms. Implementation of ‘emotion algorithms’ in artificial intelligence solutions may in some cases be feasible only *in silico* and for certain problems. In this vein, computer and decision scientists discover solutions to optimize learning and decision making that may not be possible for biological agents that are strictly constrained by limited time and energy, high uncertainty environments, and constant threats to existence (Montague 2006; Bach and Dayan 2017; Huys and Renz, 2017). Here, it must be assumed that evolution has shaped the mechanisms underlying emotion-related modifications to behaviors and associated subjective experience. Thus, empirical work (in conjunction with theory) is necessary to discover how the human brain solves the decision-making problem and generates associated subjective experiences. Computational neuroscientists trying to discover how emotions are generated are beginning to investigate how neuromodulatory systems (e.g., dopaminergic, serotonergic, and noradrenergic neurons) drive not only learning and arousal states, but also affective dynamics, mood, and emotion regulation (Doya 2002; Etkin et al., 2015; Eldar et al, 2016; Huys and Renz, 2017; Bach and Dayan 2017).

One of the first demonstrations that reward prediction errors modulate self-reports about subjective feelings came from work investigating social exchange (Xiang et al, 2013). In this work, Xiang and colleagues had participants play the ultimatum game repeatedly, but with different partners. In the ultimatum game, a proposer is given a set amount of money and propose a split of the money between themselves and a partner. The partner then has the choice of accepting the split or rejecting the offer, knowing that a rejection means that both players will receive nothing (Fehr and Gächter, 2002; Camerer, 2003). In Xiang et al., the participants were the partners receiving the offered splits and the proposers were computer agents programmed to give similar distributions of offers and to shift their proposals (as a group) halfway through the task. In this manner, participants received offers that at first may be low or high, but common to the group and a ‘social norm’ learned. Deviations about this norm generated prediction errors about the expected offer value, which were exaggerated when ‘the group’ shifted its behavior mid-way through the task. Participants playing this task were instructed that they would be playing with partners that were all different and independent. Occasionally, participants were probed about ‘how they felt’ about the most recent offer. Xiang et al., report that subjective feelings correlated with the norm prediction error and that the norm prediction error correlated with fMRI measured BOLD responses in the orbital frontal cortex (Xiang et al. 2013). Additionally, BOLD responses to the norm prediction error were observed in the dorsal and ventral striatum, and bilateral anterior insula.

Eldar and colleagues (Eldar et al., 2016) review recent applications of a reinforcement learning framework and gambling tasks to investigate affective dynamics and mood and hypothesize how the proposed computational models may be used to better understand mood disorders. Two key studies form the basis of their argument (Rutledge et al., 2014 and Eldar and Niv, 2015). Rutledge and colleagues (2014) investigated emotional reactivity in response to a probabilistic reward task that did not require participants to estimate the expected values of choices presented through learning. Expected values could be calculated on the spot (per trial) with complete information about the risks associated with either a gamble or an alternative ‘sure bet’. They demonstrated that dynamic changes in “momentary happiness” were explained by a non-linear combination of expected values and associated reward prediction errors over a short history of recent trials. Further, they used fMRI to show that BOLD responses in the striatum tracked the same task variables that predicted subjective happiness ratings. Together, these findings strongly implicate a role for parameters estimated using computational reinforcement learning theory (e.g. expected value and reward prediction errors) and the dopaminergic system (i.e., dorsal and ventral striatal activation to reward prediction errors). Importantly, striatal BOLD imaging signals that track computed reward prediction errors are only circumstantial evidence that these signals are in fact delivered by dopamine neuron activity – BOLD imaging tracks physiological activity that consumes oxygen in the blood and cannot alone distinguish activity specifically driven by dopamine release.

Eldar and Niv (2015) used a computational reinforcement learning framework to investigate a hypothesized bidirectional interaction between the ‘evaluation of outcomes’ and ‘changes in emotional state’, and how the latter impacts the evaluation of future outcomes. Participants played a series of probabilistic slot machine games before and after a “wheel-of-fortune” draw that resulted in a surprising large magnitude outcome. The impact of the wheel-of-fortune draw went as one might expect – improved mood for a big win and decreased mood for big losses, but the impact of mood was also measured on the outcomes of the subsequent smaller outcome slot machine games. They showed that large magnitude outcomes on the wheel-of-fortune game colored the outcomes of the subsequent slot machine games. Again, a role for dopamine was implicated by the demonstration of striatal BOLD responses to reward prediction errors following wins in the slot machine games. Interestingly, the impact of the wheel-of-fortune outcome on striatal responses to the slot machine game outcomes differentiated those participants who had an increased hypomanic personality. The authors then demonstrate within a reinforcement learning framework that they could account for and predict the behavioral observation that mood instability was associated with a positive feedback loop where big changes in mood altered the perception of outcomes in subsequent slot machine games. Together the results of their study implicate dopaminergic learning signals in modulating dynamic changes in affect, a process that may become destabilized in people with increased mood instability.

Another study that has embraced the reinforcement learning framework to investigate the impact of emotional outcomes on future decisions was performed by Katahira and colleagues (2015). Many studies have used monetary incentives to demonstrate the interaction of estimates of expected value, reward prediction errors, and human decision-making. The advantage of monetary incentives is the inherent quantitative nature of the reinforcer, so incorporating money as the reward in the reinforcement learning framework is relatively straightforward. Katahira and colleagues tackled an important problem in a creative way. They asked whether decision-making under uncertainty where emotional outcomes (that are inherently subjective) could be modeled and understood in the quantitative reinforcement learning framework. Here they employed a probabilistic “reward and punishment” task where instead of money, participants received feedback in the form of subjectively pleasant or unpleasant pictures, respectively. Images were drawn from the international affective picture system data base (Lang et al., 2007). While the categories of pleasant, neutral, or unpleasant images could be estimated *a priori* from the IAPS database, it is unclear what the motivational value of each image would be per participant. Typically, in reinforcement learning modeling, the learning rate (*α*) and future discount rate (*γ*) are estimated as free parameters while fitting the models to participant behavior. Katahira and colleagues allowed the ‘motivational value’ of each image category to also be a free parameter and could thus estimate from behavior a quantity that would otherwise be a private subjective value. They compared behavioral and brain responses fit to reinforcement learning models for this task to a monetarily incentivized task modeled in a similar manner and demonstrated both appetitive and aversive prediction errors in the striatum for both the emotionally evocative task and the monetarily incentivized task. Other co-activated regions included bilateral insula and precuneus.

Model-based reinforcement learning algorithms have been used to explore how emotion regulation via (re)appraisal processes may be implemented in neural systems (Etkin et al., 2015; Huys and Renz, 2017). Model-based (versus model-free) reinforcement learning refers to an approach wherein the decision-making agent uses a model of the environment to speed learning. If a model of the environment is available, an agent can use it to simulate experiences and significantly enhance the rate of learning compared to solely experience-dependent (i.e., model-free) algorithms (Sutton and Barto, 1998; Sutton and Barto, 2018). There is plenty of evidence that suggests humans (and other biological agents) use such models in their decision-making. Yet, the physical manifestation in neural systems of these representations and simulated experiences remains up for clarification. Nevertheless, model-based algorithms serve as an excellent framework to investigate mechanisms supporting emotion regulation or control. Etkin and colleagues (Etkin et al., 2015) have proposed the use of model-based reinforcement learning to unify diverse findings about the neural structures involved in the spectrum of emotion regulation strategies. In their framework, emotions are generated as part of the typical value-based decision-making process, whereas regulation of these emotions are thought of as meta to the generation process: ‘actions’ taken by the emotion regulation system are themselves ‘decided’ upon within a value-based decision-making process that adjudicates between the value of different state-action pairs representing emotion states and associated behavioral repertoires. They argue for a spectrum of model-based to model-free strategies for emotion regulation and suggest key differences in neural network dynamics that support each strategy. In a similar manner, Huys and Renz argue for the need of meta-reasoning over a model-based valuation framework to account for the nature of emotional appraisals and the flexibility of emotional responses.

The strengths of using a computational reinforcement learning framework to investigate mechanisms supporting affective dynamics include the formal nature of the description of its algorithms (Bach and Dayan, 2017; Huys and Renz, 2017) and the flexibility to explore variations on the core theme (Moerland et al., 2018). The exploration of methods to incorporate ‘emotion computations’ into reinforcement learning in artificial intelligence research (e.g., Moerland et al., 2018) provides a likely never-ending stream of ‘thought experiments’ that may or may not be relevant to the biology of human emotions. Accordingly, research that constrains the space of possible solutions using empirical results is necessary.

### Limitations of ‘dopamine/TD-reward’ -centric models

Over the last two and a half decades, the success in applying computational reinforcement learning algorithms to elucidate the neurobiological basis of human (and non-human animal) decision-making has been fueled by the singular demonstration that dopamine neurons encode the temporal difference reward prediction error (Figure 1 and equation 4; Montague et al., 1996; Schultz et al., 1997). The empirical results discussed above that relate reinforcement learning to human affective dynamics all have the reward prediction error and dopamine’s hypothesized role at their core. This core concept serves as an anchor in these early days, but there are significant challenges to this construct that must be overcome. Dopamine neurons signal temporal difference reward prediction errors by emitting a burst (an increase in firing rate for positive reward prediction error), pause (halt in firing rate for negative reward prediction error), or no change (reward prediction error equals zero) in activity (Figure 1, Montague et al., 1996; Schultz et al., 1997; Watabe-Uchida, et al., 2017). This asymmetry poses a problem for encoding aversive input – a missed reward and a loss of any magnitude would be treated the same: a pause in dopamine neuron activity. This means that a dopamine neuron-centric system (acting in this manner) cannot parametrically encode aversive prediction errors. Likewise, theories of emotion that rely only on the dopaminergic system do not appear to be able to account for subjective feelings in the negative domain, but instead focus largely on the positive dimensions like pleasure (Kringlebach and Berridge, 2017).

Further, there is now a significant body of literature that suggests that dopaminergic signaling may be more complex (Bromberg-Martin et al., 2010; and Lammel et al., 2014; Watabe-Uchida et al., 2017). Electrophysiological (and more recently optical) recordings of putative dopamine neurons in the VTA and SN (in nonhuman primates and rodents) during a variety of behavioral task demands provide evidence that dopamine neurons respond to a wide range of positive, negative, and neutral stimuli. This may not be surprising given the significant heterogeneity in the molecular, cellular, and neuroanatomical characteristics that differentiate dopamine neurons (Watabe-Uchida, et al., 2012; Lammel et al., 2014; Beier et al., 2015; Lerner et al 2015; Gantz et al., 2018).

As such, a large number of signals have been proposed to be encoded by dopamine neurons. One parsimonious way to frame these signals is to split them into ‘positive’ (or at least neutral) and ‘negative’ valence categories. We start with the positive or neutral valence category of signals as these responses may be seen as an extension of the foundational notion that dopamine neurons encode TD-error signals that, as has been discussed, may be associated with primary rewards or also predictors (e.g., context and cues) of those rewards (Montague et al 1996; Schultz et al., 1997). Then we will review and discuss the more controversial implications of dopamine neuron encoding of aversive or negatively valent stimuli.

### Dopaminergic response to novelty and surprise ....

Some of the earlier evidence that suggested that dopamine neurons encoded more than just a TD reward prediction error came from neural recordings in non-human primates (reviewed in Bromberg-Martin et al, 2010). In these experiments, it was shown that putative (electrophysiologically characterized) dopamine neurons responded in a brief burst in activity to surprising sensory events. These signals were diminished if the sensory stimulus became predictable and were absent when the animal was asleep (Takikawa et al., 2004; Strecker and Jacobs, 1985; Steinfels et al., 1983). These signals also appeared to be sensitive to the attentional demands of competing tasks, in that the ‘alerting response’ (following Bromberg-Martin et al, 2010) in dopamine neurons to unanticipated sensory stimuli could only be elicited when the animal was in a passive resting state, not engaged in a more highly-valued, goal-directed activity (Strecker and Jacobs 1985; Horvitz et al., 1997; Horvitz, 2002).

This idea that a subset of dopamine neurons respond to surprising stimuli or novelty (Ljungberg et al., 1992) has been supported by more recent work in rodents (Menegas et al., 2017). Here, Watabe-Uchida and colleagues used fiber-fluorometry to monitor dopamine axons across the striatum. They observed differences in the dopaminergic response to novel cues when they compared responses in the ventral striatum and the posterior tail of the striatum. Importantly, these responses were monitored in mice performing a classical conditioning task where the reward prediction error hypothesis could be tested. Dopamine responses in the posterior tail of the striatum responded strongly to novel cues, whereas the dopamine response in the ventral striatum did not – at least not until the novel cue was reliably paired with a reward. Like the data generated in nonhuman primates, the dopamine responses (in the posterior tail of the striatum) to novel cues diminished with experience, but these responses also occurred tied to a variety of stimuli including rewarding, aversive, and neutral sensory stimuli (Menegas et al., 2017). Additionally, Menegas and colleagues were also able to show two important controls for these responses within the same experimental paradigm: 1) specificity of dopamine neuron activity through genetic control over the reporter, but also 2) that reward prediction errors could explain the activity observed in the ventral striatum-projecting dopamine neurons, but could not explain the activity in the posterior tail of the striatum.

Interestingly, it has been reported that dopamine neurons in the VTA can promote wakefulness and arousal (Eban-Rothschild et al, 2016; Taylor et al., 2016), and that a class of putative dopamine neurons residing in the dorsal raphe are activated by salient stimuli and, in so doing, also promote arousal and wakefulness (Lu et al., 2006; Cho et al., 2017). These latter cells are atypical in that most investigations of dopaminergic activity focus on the main populations of dopamine neurons found in the VTA and SN.

### Dopaminergic responses regarding context and information ....

New, potentially informative sensory stimuli is one way to characterize “novel” or “surprising” cues. Along these lines, it has also been shown that dopamine neurons will respond as though they detect changes in the sensory features of rewards (Takahashi et al., 2017), can be modulated by changes in context (Nakahara et al., 2004), and can respond preferentially to sensory signals that provide advance information (Bromberg-Martin and Hikosaka 2009).

Schoenbaum and colleagues (Takahashi et al., 2017) recorded single-unit activity of putative dopaminergic neurons in rats during an odor-guided choice task. In this task, odors signaled the availability of vanilla or chocolate milk of varying quantities. They were able to demonstrate that the rats showed no distinguishable preference for chocolate or vanilla flavoring, but did track the quantitative value (quantity of milk) delivered. The recorded dopamine neurons tracked the prediction error of the value association (reward prediction errors over the quantity of milk delivered). But also, notably, when the flavor of the reward delivered was different from what was expected (vanilla instead of chocolate, or vice-versa), dopamine neurons showed an additional burst in activity. This was true even when the quantity of milk was the same as expected. The authors interpret this as a kind of prediction error over sensory features, which is consistent with the idea that dopamine neurons encode surprising sensory signals – here, the flavor of the milk. These experiments nicely control the expectation of sensory stimulation while modulating information in a surprising way about the reward delivered. Such additional information may not be directly related to the value of what was ingested, but may be used as an informative signal that the context of the behavioral task may have changed.

Hikosaka and colleagues (Nakahara et al., 2004) have demonstrated the ability of dopamine neurons to track reward prediction errors about informative cues (Bromberg-Martin and Hikosaka, 2009), and also shown dopaminergic activity that is best explained by models that also account for contextual changes (Nakahara et al., 2004). Nakahara et al. (2004) used electrophysiological recordings of putative dopamine neurons in non-human primates, a classical conditioning task, a context-dependent conditioning task, and computer modeling to demonstrate that at least two classes of dopamine neurons could be identified: dopamine neurons that track classic temporal difference reward prediction errors and dopamine neurons that detect context-dependent reward prediction errors.

Bromberg-Martin and Hikosaka (2009) used a choice task to demonstrate that macaques preferentially choose options that lead to advance information about the magnitude of reward to be delivered at a later time. Not only did monkeys display this behavioral preference, but dopamine neurons that showed reward prediction error responses to reward delivery and predictive cues also showed responses that increased when predictive advanced information was provided.

Together, these experiments highlight a strong connection between dopamine neuron activity and surprising information and the importance of context. New information or surprising changes in the environment may have intrinsic value to a system that is geared towards learning statistical structure about appetitive pro-survival events. New information or a surprising change in the context of delivered signals would be important to alert to and may simultaneously be treated as potentially rewarding and potentially punishing. In this sense, one might expect to also see an aversive expectation error response signaled in parallel with what we are hypothesizing as a dopaminergic appetitive expectation error reported above. These responses would not necessarily be learned ones, but rather an intrinsic valuation response to any ‘new information’ or events experienced for the first time that could be unlearned should the subsequent associations be consistently positive or negative.

### Dopaminergic responses to aversive stimuli ...?

The range of signals encoded above may generally be thought of as signals of positive valence, or at least not worse than neutral. However, there has been a significant amount of work demonstrating dopaminergic activity to aversive stimuli including noxious electric shock (Guarraci and Kapp, 1999; Young, 2004; Brischoux et al., 2009; Zweifel et al., 2011; de Jong et al., 2019), tail pinch (Mantz et al., 1989; Zweifel et al., 2011), air puff to the eye (Matsumoto and Hikosaka, 2009) or face (Cohen et al 2012), stress (Abercrombie et al., 1989; Anstrom et al., 2005), and social defeat (Cao et al., 2010). Questions that remain in all investigations of dopamine neuron function include whether the characterized neurons are in fact dopamine neurons (Ungless and Grace, 2012). Further, it is unclear whether the experimental paradigms that demonstrate dopamine neuron activity to aversive stimuli control for alternative possibilities (Tanimoto et al., 2004). For instance, any of these apparent aversive events may not be interpreted as aversive, since many of the animals may not have had any experience with these kinds of stimulation prior to the initial work of the experimenter. Thus, any of these seemingly aversive stimuli could also be interpreted by the animal as simply ‘novel’ or ‘potentially informative’ in an otherwise highly controlled environment.

### Summary and Discussion of Dopamine-centric Limitations

Altogether, these studies demonstrate that dopaminergic encoding of temporal difference reward prediction errors is only part of the story. There is clear evidence that at least a subset of dopamine neurons encode temporal difference reward prediction errors (Montague et al., 1996; Watabe-Uchida et al., 2017). Dopamine release in human striatum also encodes reward prediction errors, but also appears to integrate this term with contextual information about counterfactual outcomes (Kishida et al., 2016; Moran et al., 2018). This latter point raises questions about the mechanisms that lead to dopaminergic encoding of counterfactual prediction errors – ‘are these signals a result of modulation of dopamine-releasing terminals that natively would otherwise encode standard reward prediction errors, or are these terminals silent while counterfactual prediction errors are signaled by a different subset of dopamine-releasing neurons that innervate the same region of tissue?’. Above, we described work from the animal literature that shows that dopamine neuron activity seems to encode contextual and informative signals, and possibly even aversive signals. It is not clear whether these studies controlled for the possibility that the stimuli used to represent “non-rewarding” signals do not have appetitive associations acquired previously (e.g., due to routine animal handling procedures) or intrinsically, in that the stimuli that we associate with being neutral or aversive may be interpreted very differently by experimental animals that experience life very differently from organisms in the wild.

Another major question that is not answered by most of the studies above is the degree to which temporal difference learning is or is not at play for these dopaminergic responses. Except for the studies that confirm that dopamine neurons encode temporal difference reward prediction errors, temporal difference reinforcement learning models are not tested nor can be tested given the design of the reported experiments. Based on the data and results reported in such studies, temporal difference reinforcement learning cannot be ruled out. It may be that the reported dopamine responses are encoding temporal difference reward prediction errors in each of these cases but the experiments are not designed in a modeling-friendly manner, or that the ‘receptive field’ of the dopamine neuron responses under investigation may be tuned to a different objective function.

The foundational work in attempting to build a computational hypothesis about the generation of affective dynamics from reinforcement learning algorithms by Xiang, Rutledge, and Eldar and colleagues (Xiang et al., 2013; Rutledge et al 2014; Eldar and Niv, 2015) exposes the core role of reward prediction errors (presumably encoded by dopamine neuron activity) in dynamically modulating subjective feelings and mood. However, the models that link reward prediction errors to dynamic changes in happiness or mood do not provide an obvious mechanistic account for how these signals may be integrated in the brain, specifically in process that lead to the generation of subjective feelings.

In the case of the subjective happiness model (Rutledge et al 2014), the model demonstrates that recent expected value estimates and reward prediction errors are correlated with changes in subjective ratings. The non-linear model that they provide is descriptive in the relationship between the role of value estimates and prediction errors about these expected reward values. Notably, these model terms are shown to correspond to activity in regions of the brain known to track these signals (independent of their connection to subjective ratings), and the insula is demonstrated to become active during the introspective report. Xiang et al also implicate the striatum, orbital frontal cortex and insula (Xiang et al., 2013). These findings are an important first step in connecting neural computations and subjective experience but do not yield a clear computational hypothesis about the neural mechanisms that implement the integration of these signals into a subjective experience. Likewise, Eldar and Niv’s work (Eldar and Niv, 2015) clearly demonstrates the two-way interaction of immediately signaled reward prediction errors and a longer-term impact of mood-related computations and the impact of these interacting signals on subjective ratings. But, the models developed to account for these changes do not clearly indicate how the brain might encode such calculations.

We believe that the joint demonstrations that dopamine neurons encode temporal difference reward prediction errors and that humans and non-human model organisms use this signal to guide value guided behavior provides a solid foundation to build computational hypotheses about how the brain generates dynamically changing subjective experiences. However, to fully capitalize on this foundation, we must seek to explain disparate findings about potential alternative roles for dopamine neuron signals while building upon our current best theories about the connection between dopamine neuron signals and behavioral control (Montague et al., 1996, Schultz et a., 1997; Montague et al., 2004; Glimcher 2011).

## Extending Temporal Difference Reinforcement Learning as a functional motif that underlies affective dynamics: Valence Partitioned Reinforcement Learning

Temporal difference reinforcement learning algorithms were conceived to answer the problem of how an agent might learn about the value of different states in order to choose states that maximize ‘reward’ (Sutton and Barto, 1998). The computer agents implementing these algorithms obviously do not experience subjective reward; rather, the calculations aim to maximize a quantitative value defined by the algorithms’ objective functions. A major result in this line of research was temporal difference reinforcement learning, which provides an optimal way to learn from experience and generate optimal estimates of the value of different states, given the specified objective function. Thus, given good estimates of the expected ‘reward’ value, an agent can implement a policy to choose in a specified manner. It is intuitive to connect this objective function to reward since we psychologically associate reward with pleasure and generally seek to maximize this in our own subjective lives. However, the learning algorithm could also be applied to generate optimal estimates for other values that an organism (or agent) may need to track, such as the expected harm value or the expected cost of various acts. Given optimally derived estimates of what states lead to more harm (now and into the future), policies can be implemented to act to avoid these states. Likewise, a nervous system may need to track the expected ‘information’ value, which could then be used in policies aimed at building models or explore versus exploit tradeoff decisions (e.g., when there is ‘more than expected’ entropy in the environment a system may benefit from more exploration before exploiting based on its current estimates).

This line of thinking and theory-based proposals that hypothesize that the serotonergic system may act as an opponent signal to reward prediction errors signaled by dopamine (Daw et al., 2002; Dayan and Huys, 2008) led us to hypothesize that the temporal difference learning algorithm may be used by other systems in the brain. We remain agnostic as to which systems in the brain may provide these kinds of signals but note that other (non-reward responsive) dopamine neurons and the serotonin system are prime candidates. For now, we outline a simple extension of traditional temporal difference reinforcement learning (specifically Q-learning, Watkins 1989; Watkins and Dayan 1992) by which we partition valence into two independent dimensions: positive and negative (Montague et al., 2016).

### Classic Reward versus Valence-Partitioned Temporal Difference Reinforcement Learning

First, we describe the classic (unidimensional valence) approach to temporal difference Q-learning. Then we extend this approach by one step and partition the valence of inputs such that experienced appetitive and aversive outcomes are handled by independent systems before being integrated at the level of an overall value estimate. Again, the goal of Q-learning is to learn to act optimally (e.g., take actions in order to maximize reward) in uncertain environments based on past experience. Here, the *Quality* of an action (*a*) in a given state (*s*) is given by *Q*(*s, a*) rather than simply the value of a sate (as implied in equation 1–4) and is updated by the familiar temporal difference reward prediction error (*δ*):

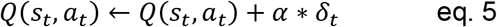

*α* is again the learning rate that determines how quickly the agent updates its expected value for the state-action pair. The temporal difference reward prediction error in Q-learning can be calculated as follows:

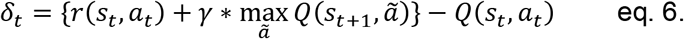

Again, this should look very familiar to eq. 4 above – now, *r*(*s*_*t*_, *a*_*t*_) is the reward collected at time *t* following the state-action pair that occurs at time *t*; *γ* is once again a discounting parameter that weights how forward looking the agent is. The value of the future state (*s*_*t*+1_) depends on the available actions (*ã*). Here, 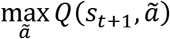 represents that value of future states when the agent chooses the action *a* out of all possible actions *ã* that maximizes the expected state-action value in the immediate future state *s*_*t*+1_. Using these Q-value estimates, the agent will enact its choice policy. The softmax policy is one policy that allows agents to make choices that balance exploiting current Q-value estimates versus exploring alternative actions so that Q-value estimates can be improved through experience. The softmax equation looks like this:

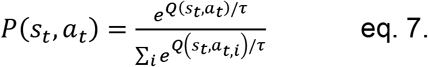

Eq 7 represents a Boltzmann distribution that specifies the probability *P*(*s*_*t*_, *a*_*t*_) that action *a*_*t*_ will be chosen in state *s*_*t*_ given the current estimates of *Q*(*s*_*t*_, *a*_*t,i*_). *a*_*t,i*_ represents each of the possible actions, indexed by *i* while in state *s*_*t*_. *τ* is a ‘temperature’ parameter that parameterizes the exploration versus exploitation trade-off: higher temperature values increase the variability of choices (weighted by the expected value *Q*(·)) while lower temperatures crystalize behavior such that actions with current maximum *Q*(·) estimates are always chosen.

To overcome limitations of the ‘dopamine/TD-reward’-centric models, we extend the traditional unidimensional valent Q-learning framework by partitioning the ‘reward’ and ‘valuation system’ into valence-specific ‘Positive’ (appetitive) and ‘Negative’ (aversive) systems. In this way, each system can be thought of as having a kind of receptive field for only appetitive or aversive inputs, respectively. We call this approach “Valence Partitioned” such that each system independently computes TD prediction errors and updates separate Q-values for positive (P) and negative (N) value estimates:

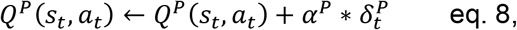

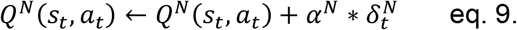

These “P” and “N” systems independently track the positive quality (*Q*^*P*^(*s*_*t*_, *a*_*t*_)) and the negative quality (*Q*^*N*^(*s*_*t*_, *a*_*t*_)) of state-action pairs, respectively. The P and N system Q-values are updated according to their own independent learning rates (*α*^*P*^, *α*^*N*^, respectively). They are updated by temporal difference prediction errors as in unidimensional Q-learning but do so only for their respective valence-specific receptive fields.

For the positive-valence *P* system, the appetitively oriented TD prediction error 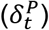 takes the form:

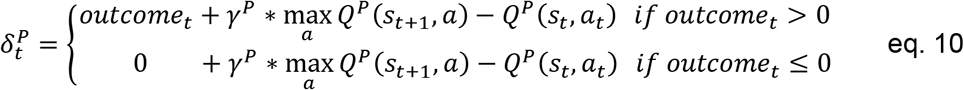

where *γ*^*P*^ is the *P* system temporal discounting parameter directly analogous to the standard unidimensional Q-learning model temporal discounting parameter.

The negative-valence *N* system similarly encodes an aversively oriented TD prediction error term 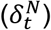:

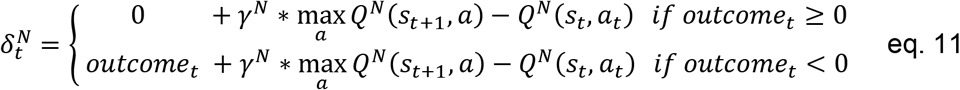

where *γ*^*N*^ is the N system temporal discounting parameter. Notably, the P system only tracks appetitive outcomes (eq. 10, *outcome*_*t*_ > 0) and the N system only tracks aversive outcomes (eq. 11, *outcome*_*t*_ < 0); otherwise, both systems ignore outcomes that are not within their receptive field (treats the opponent valence outcome as though nothing happened, eq. 10 & 11, *outcome*_*t*_ ≤ 0 or *outcome*_*t*_ ≥ 0, respectively).

Before the *P* and *N* systems’ estimates of appetitive and aversive value can be used to direct action, they must be integrated in some manner such that a policy can use these estimates to direct choice. A simple approach is to contrast them (though other schemes are possible):

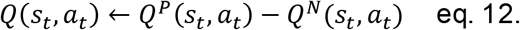

Now the output of the integrated Q-value (eq. 12) can be input in to the softmax policy equation (eq. 6; or some other specified policy) and a decision about the best action can be made.

Temporal Difference Reinforcement Learning with valence partitioning serves three purposes here. One, as a hypothetical account for how a system that is an opponent, yet complementary, to dopaminergic reward prediction error signals might behave (Daw et al., 2002; Dayan and Huys, 2008; Montague et al., 2016). Two, to serve as a hypothesis to account for observed serotonergic signals in humans (Moran et al., 2018; Bang et al., 2020). And three, as a hypothetical example of how one might extend the concept of optimal control and explore the idea that temporal difference learning algorithms are a functional motif used by multiple systems in the nervous systems of organisms that express complex adaptive behavior and behavioral control. For example, one could extend this framework such that expectations about the frequency of common versus rare signals could be optimally tracked and signaled as an ‘entropy’ prediction error that would alert the system to novel or rare signals or surprising changes in context that ought to be attended to and investigated (or avoided depending on the agent’s estimated value of new information).

Simultaneously recorded measurements of serotonin and dopamine release at sub-second temporal resolution in humans during decision-making is consistent with serotonin and dopamine acting as opponent signals (Kishida et al., 2016; Moran et al., 2018; Bang et al., 2020). In each of these reports, the measured serotonin responses are consistent with a temporal difference aversive prediction error signal and, in the case of Moran et al., 2018, are shown to anti-correlate with simultaneously measured dopamine prediction error signals that integrate reward prediction errors and counterfactual prediction errors (Kishida et al., 2016; Moran et al., 2018). Sub-second serotonin and dopamine fluctuations are also shown to anticipate actions of the participant, which may be interpreted as a signal related to behavioral control or a prediction error signal associated with anticipated outcomes – these are not necessarily mutually exclusive hypotheses.

Partitioning incoming appetitive (positive, *P*) and aversive (negative, *N*) signals would not be a challenge for biological systems and doing so would increase the dynamic range of the spectrum of valence interpretations. Behavioral results from human participants subjectively rating pictures for valence and arousal suggest that appetitive and aversive stimuli may not be unidimensional but rather independent dimensions that may be integrated in the behavioral report (Figure 3, Lang et al. 2007). For example, separate *P* and *N* systems may send prediction error signals that are integrated downstream (*P+N*) or contrasted (*P-N*) depending on the neurotransmitters and receptors that carry and receive these signals, respectively (Montague et al., 2016). The neuroanatomy of the ascending valuation systems that include dopamine, serotonin, and norepinephrine neurons are prime candidates for this kind of diverse signaling (Figure 4, Schiff and Plum 2000; Doya 2000).

**Figure 3.**
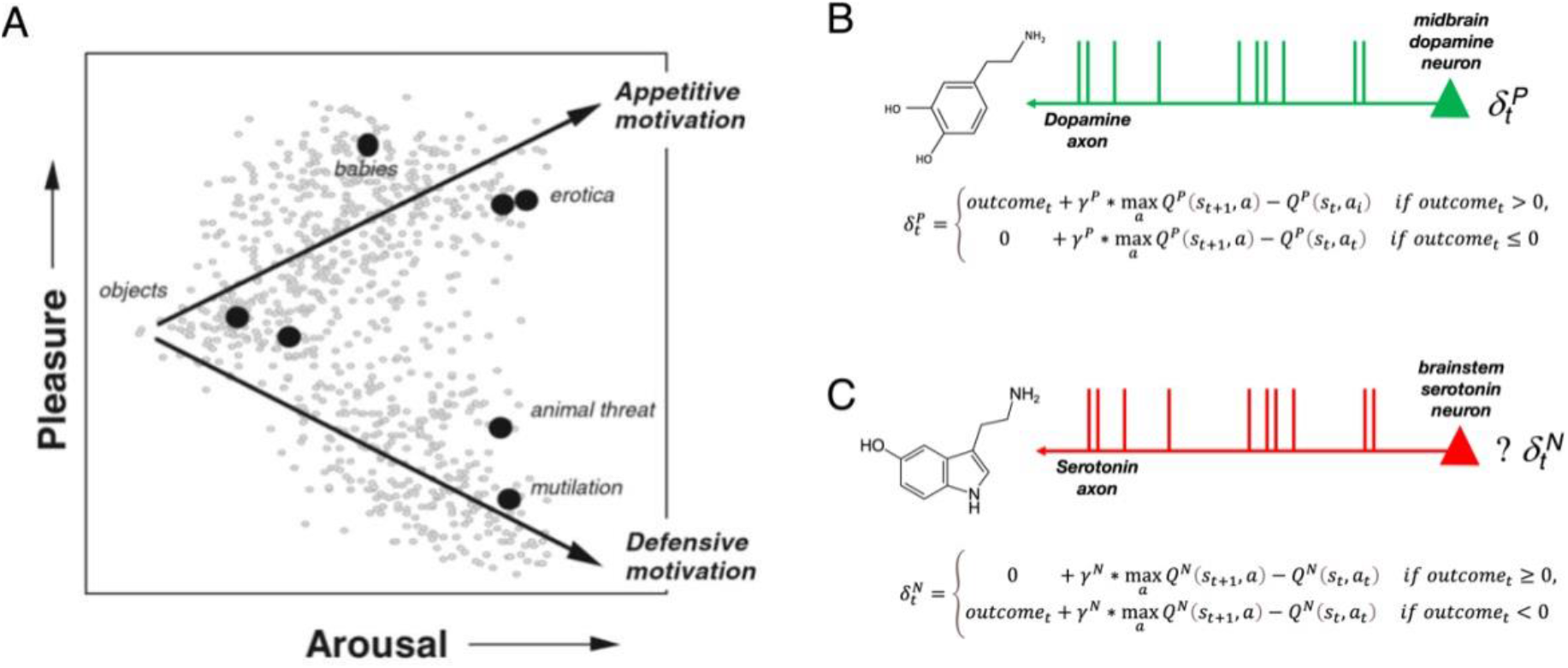
Dynamic Affect from Valence Partitioned Reinforcement Learning. **A.** The international affective pictures system (IAPS) provides a widely utilized database of emotionally evocative images that have been rated by men and women for dimensions of pleasure and arousal (and dominance) (Lang and Bradley, 2007). **B.** Valence Partitioned Reinforcement Learning (VP-RL) largely leaves dopamine neurons’ relationship to positive valence (i.e., psychologically rewarding) stimuli intact; the main difference is that the inability of this system to track variable magnitude aversive outcomes is made explicit. **C.** VP-RL posits that an additional independent system that runs in parallel but also uses TD-RL to track and estimate the expected value of aversive outcomes; we hypothesize that the serotonin system may act to track and signal aversive prediction errors 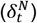, but other neuromodulatory systems can be considered as alternative hypotheses. Positive system prediction errors 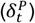, hypothesized to be delivered by dopamine are well suited to drive appetitive motivational responses as cues (e.g., images) that are predictive of positive outcomes would elicit anticipated value driven prediction errors. In a similar manner, negative system prediction errors 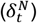 would be suited to drive defensive motivational responses. Interestingly, the independence of these systems in the VP-RL framework suggests that these signals can be integrated or contrasted by receptive systems that may in turn drive increases in arousal or enhance the discrimination of states and actio ns that result in complex valence superposition, respectively.

**Figure 4.**
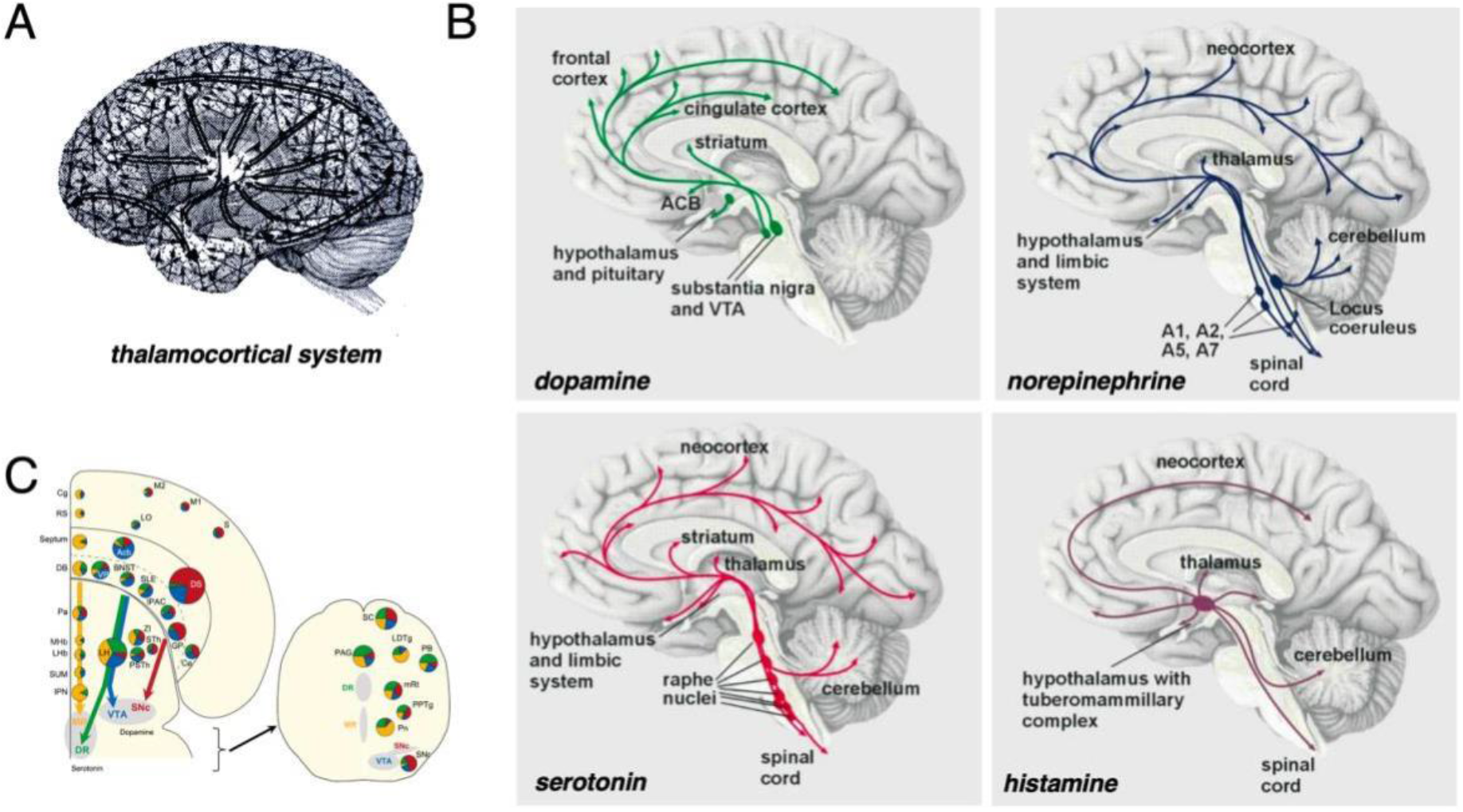
Dynamic Affective Core. The original *dynamic core* hypothesis (Edelman and Tononi 2000), similar to other neuroscientific theories of consciousness, is largely (**A**) corticothalamic centric. We propose the *dynamic affective core*, which explicitly incorporates the (**B**) ascending valuation systems (e.g., dopamine, serotonin, norepinephrine, and histamine)). Input from all over the brain is able to drive and modulate these systems wherein they provide critical neuromodulatory feedback. For example, (**C**) In the mouse, where detailed neuroanatomical tracing can be performed with the help of viral and sophisticated genetic tools, dopamine and serotonin neurons are observed to receive monosynaptic input from and send directly back signals to the amygdala, hypothalamus, thalamus, insula, cingulate cortex, striatum and many other cortical and subcortical structures (Ogawa et al., 2014). **A**. Schematic of corticocortical and thalamocortical connections that comprise the core structures that dynamically encode and represent the contents and spatiotemporal context of conscious experience (images adapted from Edelman and Tononi 2000). **B**. Schematic of ascending projections of dopamine, serotonin, norepinephrine, and histamine neurons from the midbrain and brainstem (images adapted from Fuchs and Flügge, 2004). **C**. Schematic of monosynaptic inputs to dopamine (ventral tegmental area, VTA and substantia pars compacta. SNc) and serotonin (median raphe, MR and dorsal raphe, DR) neurons in the mouse brain (images adapted from Ogawa et al., 2014). Neural structures that project to the dopamine and serotonin neurons in their respective nuclei are color coded according to their projection target. VTA: blue; SNc: red; MR: yellow; and DR: green. Pie charts indicate the proportion of neurons in that region that project to each of the respective nuclei.

## Subjective Experience and the ‘Dynamic *Affective* Core’ hypothesis

“Subjective feelings” are at the core of what humans try to describe when we communicate our experience. The contents (i.e., objects and spatio-temporal context) of our experience are part of the description, but it is the subjective feeling bound to those elements that drive our descriptions. Fundamentally, these phenomenal experiences are integral to how we perceive, navigate, and communicate about the world and consist not only of the informative content of the environmental context we find ourselves in, but critically the subjective feelings associated with it. “How we feel” from moment-to-moment impacts our mood and behavior (and vice versa) and likely involves an extensive network of dynamically interacting neural structures. Edelman and Tononi proposed the *dynamic core* as a hypothesis about how neural activity may generate conscious experiences (Tononi and Edelman, 1998; Edelman and Tononi, 2000). The *dynamic core hypothesis*, as originally stated (Edelman and Tononi, 2000), has two fundamental parts:

1. “A group of neurons can contribute directly to conscious experience only if it is part of a distributed functional cluster that, through reentrant interactions in the thalamocortical system, achieves high integration in hundreds of milliseconds.”
2. “To sustain conscious experience, it is essential that this functional cluster be highly differentiated, as indicated by high values of complexity.”

In this hypothesis, cortical-cortical and thalamo-cortical loops are fundamental to the ‘dynamic core’; timing is key as is reverberant activity such that there is wide-spread activity (throughout the cortex) that is tightly coupled in a brief window of time. Further, not just any coupled activity will do: they propose a complexity metric that is aimed at identifying specific levels of information integration and differentiation. This notion has evolved into the Integrated Information Theory of consciousness with phi (Φ), a measure of *information integration*, central to the theory (Tononi et al, 2016 for a recent review). In the original formulation of the *dynamic core hypothesis,* the ascending valuation systems that include dopaminergic, serotonergic, norepinephrinergic, and cholinergic neurons (Schiff and Plum, 2000) is noted for their likely role in *dynamically modulating behavior* in response to “external stimuli, learning and memory, emotion, and cognition” and in coupling “value and emotions” to conscious experience (Edelman and Tononi, 2000). Within the framework of the dynamic core hypothesis, it seems that the role of the ascending valuation systems is simply to shape the cortical networks based on the organism’s life experience (learning) and to drive behavioral dynamics *in response to* subjective emotional experience.

While we find the dynamic core hypothesis useful in depicting the notion of a momentary, widely distributed functional cluster of neural activity that is necessary for representing the contents of a conscious experience, we believe it lacks a critical component of subjective phenomenal experience – the emotional subjective affect that *colors* what would otherwise simply be a dry representation of integrated information collected by the sensory systems.

The original dynamic core hypothesis (Edelman and Tononi, 2000) does not include an account for the central role dopaminergic signals likely play in driving dynamic changes in subjective experience. Though the role dopamine neurons play in signaling reward prediction errors was demonstrated at the time (Montague et al, 1996; Edelman and Tononi, 2000), those findings were not discussed. Since then, the dynamic core hypothesis seems to have evolved into Integrated Information Theory where the main focus has been on developing a theoretic model that quantifies the structure of information integration that would support consciousness – whether it be biological or otherwise – and less emphasis has been placed on determining how any such a dynamic core may generate conscious experience.

### The Dynamic Affective Core Hypothesis

We propose to update the *dynamic core* hypothesis to include the ascending valuation systems and suggest that emotional circuitry and ascending neuromodulatory systems are critical components of a *dynamic affective core*. We hypothesize that this *dynamic affective core* (Figure 4) is necessary for human subjective experience. In contrast to the dynamic core hypothesis, we view the neuromodulatory systems (i.e., dopaminergic, serotoninergic, adrenergic, cholinergic, and histaminergic neurons) as a necessary component that was excluded from the original dynamic core description. Also, in contrast to the dynamic core, we hypothesize that emotional circuitry is a necessary element of the dynamic affective core that generates human subjective experience and that together the emotional circuitry and ascending neuromodulatory systems engenders the dynamic core content with the “what it is like…” aspects of subjective feelings and qualitative conscious experience. We believe that to make progress in understanding the nature of the dynamic affective core and how it generates consciousness, computationally constrained hypotheses will be necessary and will include the foundational work connecting temporal difference learning algorithms to dopamine neurons and their cortical and subcortical targets.

### Experimental Support for the Dynamic Affective Core Hypothesis

Much research has been done to elucidate neural structures that support neural representations of emotion-related behavior and subjective phenomenal experience (LeDoux, 2000; Lang and Bradley, 2010). These structures (e.g., amygdala, insula, striatum, and orbital frontal/ventromedial prefrontal cortex) and their dynamic interaction are hypothesized to be critical elements in a dynamic affective core that supports a dynamically evolving representation of continuously evolving emotional states that are in turn bound to dynamically evolving contextual sensory states. To determine the composition of these dynamic systems and their behavior in regard to affective dynamics, we turn again to computational approaches to human neuroscience.

Above, we reviewed recent research that connects reward prediction errors (putatively encoded by dopamine neuron activity) with dynamic changes in subjective feelings about social gestures (Xiang et al., 2013), subjective well-being (Rutledge et al., 2014) and mood (Eldar and Niv, 2015). It has also been shown that reward prediction errors are associated with sub-second fluctuations in dopamine release (Kishida et al., 2016; Moran et al., 2018) and aversive prediction errors are associated with sub-second serotonin release (Moran et al., 2018) in the striatum, both of which are consistent with the VP-RL hypothesis described above. In both of these studies dopamine and serotonin fluctuated in ways that were consistent with how participants’ feelings ought to have been modulated given the reward prediction error and counterfactual error signals present in the game. All together, these results are consistent with the dynamic affective core hypothesis, though significant gaps remain. Direct dopamine and serotonin measurements with the temporal resolution required are currently restricted to the striatum, though technology appears to be on the horizon that may overcome this challenge and permit real-time neurochemical detection throughout the human brain (Montague and Kishida, 2018). Dopamine and serotonin (and norepinephrine) neurons project from the midbrain and brainstem throughout the human brain (Figure 4) and whole-brain connections to the dopamine and serotonin neurons have been demonstrated (Watabe-Uchida et al., 2012; Ogawa et al., 2014). Thus, through direct and indirect pathways, dopamine neurons likely broadcast prediction error signals throughout the brain. Consistent with this idea, one ought to see evidence of reward prediction errors modulating whole networks, which would be consistent with dopamine neurons providing distributed parallel signals capable of shaping and also driving the synchronous activity of a dynamic affective core.

### Reward prediction errors modulate task specific dynamic cores

A few studies have begun to look for evidence that reward prediction errors modulate whole functional networks. These studies were performed investigating the role of reward prediction errors in instrumental reinforcement learning or associative learning tasks and did not probe associated subjective experiences. Nonetheless, these studies do demonstrate evidence that reward prediction errors modulate network level dynamics consistent with task specific dynamic cores.

Using an associative learning task with BOLD imaging, den Ouden and colleagues demonstrated that prediction errors in the striatum modulate cortical coupling (den Ouden et al, 2010). In this task, participants were required to generate motor responses indicating whether an auditory tone (high or low) was followed by one of two visual stimuli (human faces or houses). The probability of each tone being followed by each visual stimulus was changed over time. Using a Bayesian learner model, expectations (i.e., probability that a visual stimulus would occur) were estimated throughout the task and violations of those expectations generated prediction errors that parametrically modulated brain activity measured using fMRI. Non-specific prediction error signals were observed in the putamen and premotor cortex, whereas activity in the fusiform face area (FFA) was found to correlate with the probability of observing human face stimuli, and activity in the parahippocampal place area (PPA) correlated with the probability of observing houses. As such, responses in the FFA and PPA to faces and houses, respectively, appeared to be a function of how surprising each stimulus was, indicating that FFA and PPA regional activity is modulated by prediction errors over expectations of stimulus occurrence. Notably, nonlinear dynamic causal modeling was consistent with the hypothesis that prediction error signals in the putamen (a site of significant dopaminergic innervation) served to modulate functional connections from FFA and PPA to the dorsal premotor cortex. In all, these results demonstrated that regional activity in visual cortex is modulated by learning-relevant prediction error signals and that inter-regional functional connectivity within a visuomotor network (i.e., a task specific dynamic core) is modulated by prediction error signals emanating from human striatum. These findings are in line with a previous report (den Ouden et al., 2009) which used dynamic causal modeling to determine that learning-induced activity in visual cortex reflecting prediction error signals was mirrored by alterations in inter-regional functional connectivity between auditory and visual cortices, with auditory-to-visual top-down connectivity positively correlating with the prediction error-dependent regional activity in visual cortex.

BOLD activity in human striatum has also been shown to dynamically interact with distributed brain regions in visual, motor, and prefrontal cortices in a manner consistent with orchestrating valence-processing mechanisms required for reinforcement learning (Gerrarty et al., 2018). In an fMRI scanned instrumental conditioning task, participants learned which of two cues was paired with a visual stimulus. The pairings were associated probabilistically, and participants were given feedback (i.e., showing the words “correct” or “incorrect”) at the end of each trial. Choice behavior on this task was modeled such that reward prediction errors could be calculated during learning and these parametric responses could then be used to investigate changes in dynamic network connectivity using sophisticated computational tools for quantifying network dynamics (Medaglia et al., 2015). In particular, Gerrarty and colleagues were interested in investigating how a measure of “striatal flexibility” changes during reinforcement learning. Here, flexibility is a measure of dynamic network activity that indicates the degree to which a brain region functionally interacts with different brain networks over time (Bassett et al., 2011). The results demonstrated that striatal flexibility was associated with learning of cue-stimulus associations throughout the task – negatively correlated with model-derived individual-differences in learning rates, and positively correlated with individual choice temperature parameter values. Moreover, the increased striatal flexibility association with learning was determined to be implemented by increased functional coupling between nucleus accumbens and putamen with regions of visual cortex and by functional coupling between putamen and orbitofrontal and ventromedial prefrontal cortices. Together, these results suggest that regions that are highly innervated by dopaminergic inputs (e.g., nucleus accumbens, putamen, and orbitofrontal and ventromedial prefrontal cortices) are co-driven during instrumental tasks with positively- and negatively-valent feedback. This kind of dynamic network activity is consistent with our dynamic affective core hypothesis.

These studies demonstrate how the influence of putative dopaminergic reward prediction error signals may be investigated in non-invasive studies. However, direct measurement will be needed to determine whether these changes in network level representations are driven directly by dopaminergic fluctuations at key nodes or indirectly through the influence of nodes one or more synapses away within the dynamic affect core. These studies also did not investigate associated subjective experiences throughout these tasks, and future work will require experiments aimed at directly investigating the interaction of these network level dynamics and moment-to-moment changes in affect. One would expect, in these studies, that insula, amygdala, orbital frontal/ventromedial prefrontal cortex, and the striatum would be engaged as specific components of the dynamic affective core based on their previously demonstrated role in emotion and value-based decision-making behavior.

Other aspects of the *dynamic affective core hypothesis* that are not addressed in these studies is the relative lack in temporal precision and electrochemical specificity that constrains interpretation of BOLD imaging data. These studies clearly implicate regions of interest for further investigation, but BOLD imaging does not provide the temporal resolution required to observe the hypothesized affective core temporal dynamics of hundreds of milliseconds or less. Nor does fMRI provide the electrochemical specificity to discriminate the sources of neural activity that drive the blood-oxygen-level-dependent response (e.g., synaptic field potentials, somatic action potentials, or fast synaptic neurochemical signals). Studies utilizing MEG or intracranial measurements of human brain activity during conscious subjective experience may provide data with the requisite spatiotemporal resolution and – in the case of intracranial human electrochemical measurements (Montague and Kishida, 2018), the requisite neurochemical specificity – to determine the necessary and sufficient components and behavior of the dynamic affective core.

## Summary and Future Directions

Approaches using computational neuroscience to investigate *affective dynamics* are beginning to suggest methods that will enable objective investigations about the hardest problem in consciousness research. The subjective phenomenal experience of “what it is like…” to be human is not just about the contents (i.e., the objects and spatiotemporal context) of our experience. Rather, the subjective feelings or values that are also bound to these contents are what make qualitative phenomenal experience (i.e., qualia) unique to the experiencing individual. These values are likely learned (or at least modulated) through experience, become associated with states and actions that result from past states and actions, and become associated with states and actions that are predictive of future experiences as we move through space and time. Dynamic algorithms derived from artificial intelligence research (i.e., temporal difference reinforcement learning) that learn and change based on experience appear to be embodied by key neurobiological substrates and have begun to provide critical insight into how human nervous systems may encode subjective value. These models, in combination with computational approaches to characterize complex dynamic network structure, are beginning to permit the expression of quantitative hypotheses about the neural architecture that supports conscious subjective experience in humans. This, in turn, provides guidance regarding the kinds of experiments required for advances in consciousness and affective dynamics research.

We introduce the *dynamic affective core* hypothesis. It is different from the original ‘dynamic core’ hypothesis as stated by Edelman and Tononi (2000) and previous notions of a psychological affective core, which are not necessarily tied to the neural structure and dynamics that are central to the *dynamic affective core* that we describe. The dynamic affective core departs from the original notion of the *dynamic core* by specifically requiring the ascending valuation systems (i.e., dopaminergic, serotonergic, noradrenergic, and histaminergic neurons) and emotional circuitry (e.g., amygdala, insula, anterior cingulate cortex, and orbital frontal cortex) as necessary elements. The interaction of these systems is hypothesized to be fundamental to the binding of subjective feeling and affective value to the contents (i.e., objects and spatiotemporal context) represented in the cortico-thalamic systems of focus in the original dynamic core hypothesis. In this way, the dynamic affective core hypothesis may serve as a bridge connecting cognition, emotion, and the impact of ‘surprises’ on individuals emotional experience (Mellers et al., 2013). In the dynamic affective core hypothesis, we retain the notion of the functional cluster and timing aspects of the original dynamic core hypothesis; however, we posit that the role of the ascending valuation systems (like the distributed dopaminergic signal) is necessary for not only the selection of which functional clusters are active but also for creating and maintaining functional clusters that bind subjective value and emotion to the contents and context that activity in the cortico-thalamocortical networks encode. The timing of dopamine neuron activity (Figure 1) and modulations in extracellular dopamine levels (Figure 2) are consistent with the requisite timing to achieve “high integration in hundreds of milliseconds”. Also, the anatomical projections of dopamine (and other ascending valuation systems) neurons (Figure 4) make plausible parallel simultaneous signaling to cortical, thalamic, and sub-cortical structures which would allow adaptive formation of new functional clusters or dynamic modulation of existing ones. The dynamic affective core hypothesis is also significantly different from IIT (Tononi et al, 2016), which is primarily concerned with quantifying dynamic network structure (the amount of information integration and differentiation in a dynamic network) that might support conscious experience. The dynamic affective core is also distinct from a global workspace theory (Baars, 1997) or a global neuronal workspace as proposed by Dehaene and Naccache, 2001. Chiefly, like the dynamic core and the global neuronal workspace hypotheses, the dynamic affective core is grounded by neurobiological data; however, unlike the dynamic affective core hypothesis, both the dynamic core and the global neuronal workspace hypotheses appear to be extremely cortical-thalamocortical centric. Some of the functional attributes of the ‘workspace neurons’ in the global neuronal workspace appear consistent with the long range and diffuse projections of dopamine, serotonin, and norepinephrine neurons, but to our knowledge these neurons are not explicitly discussed as likely candidates (Dehaene and Naccache, 2001); further, dopamine, serotonin and norepinephrine neurons appear to have functional attributes that either go beyond the role of ‘workspace neurons’ to the point that they may be inconsistent with this notion. Our dynamic affective core hypothesis does not reject the critical role of cortical-thalamocortical networks and reentrant activity, it does require that the dopaminergic system (and other valuation systems) be considered as critical components that co-activate, shape, and coordinate network activity underlying affective dynamics and associated subjective feelings.

The role (specifically) of the dopaminergic system in the dynamic affective core implies a role for temporal difference learning algorithms and dynamic modulation of the dynamic affective core though the delivery and representation of the computational signals described in this framework. We want to make explicit this connection in our hypothesis. The dynamic affective core hypothesis requires computational depictions of dynamic network structure in order to move the work forward. We propose a first step in extending the dynamic affective computational framework with our valence-partitioned reinforcement learning (VP-RL) model. VP-RL describes one way that the dynamic affective core may be modulated independently by positive and negative input, which are expected to be able to also occur independently in nature. VP-RL prediction error signals about reward and punishment are consistent with simultaneously recorded sub-second data of dopamine and serotonin in humans (Kishida et al., 2016; Moran et al., 2018) and with the non-linear relationship of pleasure and arousal in consciously rated IAPS images (Figure 3). The neurocomputational framework put forward in the dynamic affective core hypothesis may also be utilized to investigate the neurobiology underlying emotion behavior observed in reaction to ‘surprising’ information in real world settings (Mellers et al., 1997 and 2013; Bhatia et al., 2019). At the behavioral level, errors in predicted outcomes seem to drive the strongest emotional reactions (Mellers et al., 1997; Villano et al., 2020). Other ideas that are likely to inform models about subjective experience and the neurobiological dynamics that support it are likely to come from research exploring the role of various reinforcement learning model elements like variations in model-based implementations or off-policy learning strategies (Doya, 2000; Montague et al., 2006; Montague et al., 2016; Bach and Dayan, 2017; Huys and Renz, 2017;) and how they interact with dynamic functional networks. To the latter, we argue for the use of computational models and descriptions of network architecture (Bassett et al., 2011; Medaglia et al., 2015). As research connecting these areas emerges, we speculate the need for mathematically depicted network structures to describe the details of the structure and function of the dynamic affective core and its relationship to conscious subjective experience.

We focus much of our review on the neurobiology of dopaminergic signals and their connection to temporal difference reward prediction errors. These neurons and their impact on behavioral control and learning have a rich literature that is grounded on the initial discovery of Montague, Dayan, and Sejkowski (1996) that dopamine neurons encode temporal difference reward prediction errors. Other systems that are certainly involved in dynamic affective processing and consciousness include the serotonergic, noradrenergic, and cholinergic and histaminergic systems (Schiff and Plum, 2000). Attempts to model these other systems has yet to yield a clear picture, but current evidence strongly suggests a role for serotonin in aversive processing (Dayan and Huys 2008; Cools et. al., 2011; Rygula et al., 2015; Moran et al., 2018; Bang et al, 2020; Doya, et al., 2021) and norepinephrine and cholinergic signals in arousal and attention (Schiff and Plum, 2000). Our valence-partitioned reinforcement learning model is consistent with dopamine and serotonin acting as opponent (orthogonal) systems that signal appetitive and aversive prediction errors, respectively and independently (also see Montague et al., 2016). Moran and colleagues observed that dopamine and serotonin release appear to encode these signals in the striatum simultaneously and in a colocalized manner in humans, which suggests that downstream dopamine and serotonin receptors may integrate or contrast these signals to give rise to a spectrum of neural interpretations. Notably, an integration of appetitive and aversive prediction errors is consistent with a saliency signal that may directly or indirectly engage arousal systems and direct attention towards relevant environmental features; contrasting these signals would increase the signal to noise in systems gauging whether to increase or decrease appetitive versus aversive representations (Montague and Kishida et al., 2016). Clearly, more work is needed to elucidate what dopamine, serotonin, norepinephrine, acetylcholine, and histamine release encode in the human brain and what role these signals play in affecting the hypothesized dynamic affective core.

We hope it is apparent that we should no longer ignore the investigation of behavior directly associated with subjective human experience (i.e., self-report). Quantifying subjective experience is challenging. Combining questions like, “How much pain do you feel?” with a visual analogue scale *is* the gold standard for assessing pain in clinical settings and really the only way to directly capture how a person feels in the moment. What is good about the visual analogue scale (and related measures) is that it reliably assesses subjective feeling and is a semi-quantitative and reproducible choice behavior. The latter points allow researchers to integrate subjective self-report ratings with the computational models (e.g., reinforcement learning framework) and model the answers as value-based decisions dependent on states and available actions. This approach, in combination with computational depictions of network structure and dynamics, will allow the field to translate reports about subjective feelings into an empirical and mathematical framework capable of precise hypothesis testing (i.e., computational modeling) analogous to what several groups have begun to do (Xiang et al., 2013; Rutledge et al.,2014; Eldar and Niv, 2015).

We propose the *dynamic affective core* hypothesis to be tested within the domain of computational human neuroscience to tackle “the hard problem” of consciousness. “The hard problem” of consciousness originally stated by Chalmers (1996) has stood as a major barrier in scientific progress in consciousness research and the subjective aspect of affective dynamics. We take the position that prescientific descriptions of phenomena always appear as “hard problems” given that the state of scientific knowledge at the time of such distinctions are not up to the task of making the problem appear ‘easy’ (Churchland 2005). We are in a time where there is so much neuroscientific knowledge that it may seem inconceivable that there are natural neural phenomena yet to be unraveled, but human consciousness stands as a definitive example. Our challenge is to convert consciousness research from a philosophically hard problem as it seems to be to one that has some scientific traction. This can only be achieved if we tackle it as an ‘observable’ phenomenon even if such observations are initially reliant on simple behavioral reports. The way through will be to constrain our hypotheses with mathematical models that relate brain and behavior such that errors and assumptions can be clearly observed and corrected. We introduce the dynamic affective core hypothesis as an idea that can take us further down this path and look to an integration of the computational neuroscience of decision-making and network dynamics to lead the way forward.

## Acknowledgements

This work was supported by funding from the NIH. KTK was supported by the NIH-NIDA (R01-DA048096), NIH-NIMH (R01-MH121099), NIH-NINDS (R01-NS092701), and NIH-NCATS (KL2TR001421). PS was supported by NIH-NIDA (T32-DA041349 and F31-DA053174).

## Notes

### Competing Interest Statement

The authors have declared no competing interest.

